# A non-parametric Bayesian prior for causal inference of auditory streaming

**DOI:** 10.1101/139188

**Authors:** Tim Yates, Nathanael Larigaldie, Ulrik R. Beierholm

## Abstract

Human perceptual grouping of sequential auditory cues has traditionally been modeled using a mechanistic approach. The problem however is essentially one of source inference – a problem that has recently been tackled using statistical Bayesian models in visual and auditory-visual modalities. Usually the models are restricted to performing inference over just one or two possible sources, but human perceptual systems have to deal with much more complex scenarios. To characterize human perception we have developed a Bayesian inference model that allows an unlimited number of signal sources to be considered: it is general enough to allow any discrete sequential cues, from any modality. The model uses a non-parametric prior, hence increased complexity of the signal does not necessitate more parameters. The model not only determines the most likely number of sources, but also specifies the source that each signal is associated with. The model gives an excellent fit to data from an auditory stream segregation experiment in which the pitch and presentation rate of pure tones determined the perceived number of sources.

## Introduction

Ambiguity in perceptual systems is a blight for inference. When we hear two sounds sequentially, we may infer that they came from two different sources, A and B, or the same source repeated. A third sound is heard – are the sources AAA, AAB, ABA, ABB or ABC? By the time four, five and six sounds are heard the number of combinations reaches 15, 52, 858. The ambiguity breeds to generate a combinatorial explosion, and yet the human auditory system is able to reliably allocate multiple sources of sound in complex, real world situations. Features of the signal are consistently associated with different sources, allowing us to keep track of a speaker’s voice and the wail of an ambulance siren, separate from the noise of background traffic and falling rain.

For several decades, the human ability to segregate sequential sounds into streams corresponding to sources has been investigated using simple sequences of either pure tones or more complex sounds (reviewed in (B. C. J. Moore & Gockel, 2012)). The time interval between tones, their pitch difference and the duration of a sequence are among the factors that play an important role (Anstis & Saida, 1985; Bregman & Campbell, 1971; van Noorden, 1975): explanations of how the factors are used based on principles such as Gestalt laws and Occam’s razor have been incorporated into the sophisticated conceptual model of Bregman (Bregman, 1994). Descriptive models based on peripheral excitation (Beauvois & Meddis, 1997), coherence of coupled oscillators (Wang, 1996) and cortical streaming modules (McCabe & Denham, 1997) provide mechanisms to estimate the number of streams, but do not specify which sound is associated with which source. While some of the models are expandable to allow more sources to be inferred, it is not known if they would cope with the combinatorial explosion. Furthermore, Moore & Gockel (B. Moore & Gockel, 2002) conclude from an extensive review of the literature that any sufficiently salient factor can induce stream segregation. This indicates that a more general model of inference is needed, that can incorporate any auditory perceptual cue and multiple sounds with different sources.

If ambiguity is a blight for inference, regularities in natural signals are the cure. Not all combinations of signal sources are equally likely – when perceptual systems generate a model of the world, we assume that they infer the most likely interpretation because the perceptual systems are optimized to the statistics of natural signals (Barlow, 1961; McDermott & Simoncelli, 2011). Bayesian inference has had considerable success in modeling many visual and multi-sensory percepts as a generative, probabilistic process (Shams, et al. 2005; Weiss et al. 2002). Despite these successes, and the increasing evidence for the importance of predictability for auditory perception (for a review see Bendixen, 2014), we still have no general, principled model of how the auditory system solves the source inference problem.

A Bayesian approach to auditory stream segregation has been used to model the dynamics of perceptual bistability (Lee & Habibi, 2009) but assumes that only two percepts are possible. Turner (2010) has developed methods of analyzing statistics of sounds based on Bayesian inference, and constructed a model to synthesize realistic auditory textures. Promisingly, inference in the model can qualitatively replicate many known auditory grouping rules.

In our model the probability of many alternative stream configurations (given the input signal) are calculated and the percept generated corresponds to the most probable configuration. The probabilities are calculated using Bayes’ rule to combine the likelihood of generating a signal given a postulated stream configuration, with the prior probability of sounds being associated with different sources. The likelihood and prior probability distributions are iteratively updated in a principled manner as information accumulates. The forms of the distributions are presumably optimized to natural signal statistics: the likelihood distribution we use is based on considerations of the physical limitations of oscillators. However, the framework of the model allows formulations of multiple explanatory factors, such as those determined by Bregman (1994) from psychophysics experiments, to be simply incorporated in the distributions. Furthermore, while the current study uses simple pure tones (replicating work by Bregman), the framework allows more complex cues from audition and other modalities to be used as long as their perceptual difference can be quantified.

## Human inference model

Pure tones are the indivisible atoms of input to the model – each being assigned to just one sound source, or stream. Inspired by work done on non-parametric priors (Froyen, Feldman, & Singh, 2015; Orbanz & Teh, 2010; Wood, Goldwater, & Black, 2006) we assume the existence of an infinite number of potential sources, leading to a sequence of tones with pitch *f_1_*, *f_2_*…, onset time 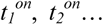 and an offset time, 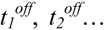 and the sound sources/streams that generated the tones are denoted by positive integers *S_1_, S_2_*… We rename the sources when necessary so that the first tone heard will always be generated by source 1 (i.e. *S_1_* = *1*), and a subsequent tone, *S_n_* can be associated with source *1:max(S_1_…S_n-1_)*+*1*.

### Generative model

Given a source *S_i_* we assume that the frequency of tone *i* is governed by physical constraints and statistical regularities of the source. If two sounds *f_1_* and *f_2_* with frequencies *F_1_* and *F_2_* are produced by the same source, the pitch cannot change at an infinitely fast rate: to make an oscillator change its frequency discontinuously would require an infinite impulse of energy. We assume that, all things being equal, a pure tone sound source is most likely to continue oscillating at the same frequency as it has in the past, and the probability of it changing at a rate Δ*F*/Δ*t* will decrease as Δ*F*/Δ*t* increases. More specifically we assume a normal probability distribution:

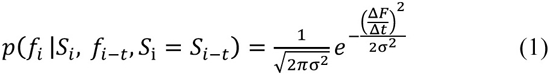

where *σ* is a constant. We here assume that the observer has a perfect noise free access to the generated frequency.

### Inference

The task of the observer is to infer the sources generating each of the tones, i.e. to find the *S_1_ S_2_ S_3_*… that maximize *p(S_1_ S_2_ S_3_… | f_1_ f_2_ f_3_…)*, as illustrated in figure 1. As an example we use a sequence of three tones *f_1_ f_2_ f_3_*, for which the observer wishes to infer the likely sources *S_1_ S_2_ S_3_*. Thus the probability *p(S_1_ S_2_ S_3_ | f_1_ f_2_ f_3_)* that a sequence of three tones was generated by sources *S_1_ S_2_ S_3_*, has to be calculated over the five combinations: [*S_1=_1, S_2=_1, S_3=_1*], [*S_1=_1, S_2=_1, S_3=_2*], [*S_1=_1, S_2=_2, S_3=_1*], [*S_1=_1, S_2=_2, S_3=_2*], [*S_1=_1, S_2=_2, S_3=_3*] corresponding to the five unique configurations of sources generating three sounds. Note that the first source is always assigned the value 1, the next different source is assigned 2, etc.. Bayes’ rule relates each conditional probability (the posterior distribution) to the likelihood *p(f_1_ f_2_ f_3_| S_1_ S_2_ S_3_)* of each configuration of sound sources generating the sequence of tones, by

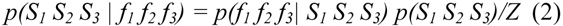

where *Z* is a normalization constant, and *p(S_1_ S_2_ S_3_)* is the prior probability of the particular configuration of sound sources, regardless of the frequency, etc. of the tones

**Figure 1:**
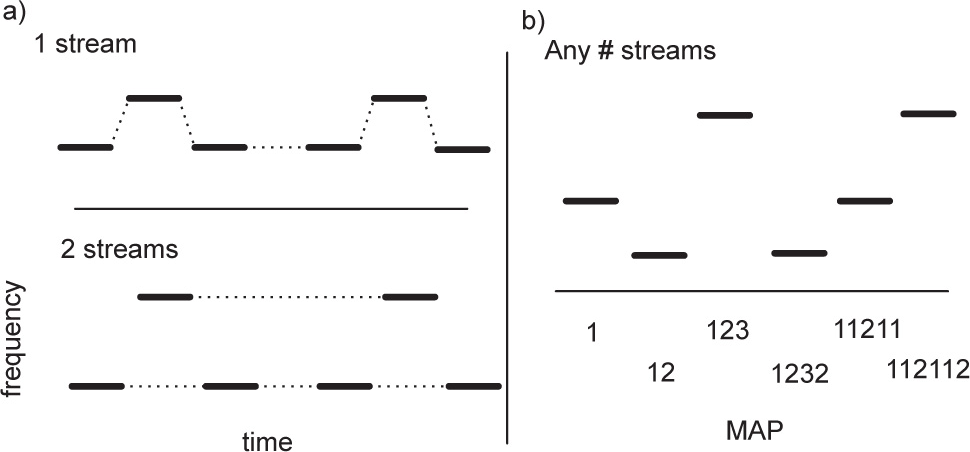
a) Example of the integration or segregation of tones, either as 1 stream or 2 streams. b) Example of the condition [3 1 9 1 3 9] from Exp. 2 (top) and the model’s sequential maximum a posteriori assignment of tones within a stream (bottom). As each tone arrives the model reassigns the entire set of tones to streams (1->12->123 etc.).

Assuming conditional independence of the tones and tone-source causality, this can be rewritten as

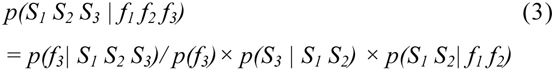

The final term is the posterior generated from the first two tones. The latter two terms can be considered together as the prior for the third source, allowing us to use an iterative approach to the inference. After each tone we grow the tree of possible source sequence (e.g. *11* → *111* and *112*), by multiplying the previous posterior *p(S_1_ S_2_| f_1_ f_2_)* with two terms; the likelihood *p(f_3_| S_1_ S_2_ S_3_)* and a prior for how likely the next ‘branch’ is, *p(S_3_ | S_1_ S_2_)*.

We now consider how to determine the likelihood and prior probabilities. The first source can only be associated with one source, so *p(S_1_=1) = 1*. The principle of Occam’s razor would suggest that *p(S_1_=1, S_2_=1) > p(S_1_=1, S_2_=2)*, i.e. if we haven’t heard any of the sounds, the most probable acoustic scene is the simplest one: all sounds come from the same source. The value of *p(S_1_=1, S_2_=1)* for an individual can be determined from fitting their data, and the value *p(S_1_=1, S_2_=2)* is simply *1– p(S_1_=1, S_2_=1).* The values may depend on factors such as the environment, which are not considered in the model: natural signal statistics may provide guidance for how the prior probabilities are assigned. For successive sources, we use the probability given by a Chinese restaurant process (CRP) (Aldous, 1985), which can be considered as an extension of Occam’s rule:

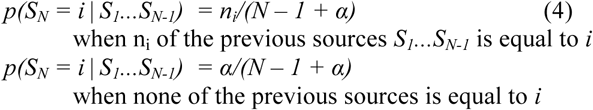

where *N* is the total number of sounds heard.

Regarding the likelihood function, the observer assumes the generative probability *p(f_i_ | S_i_, f_i-t_, S_i_ = S_i-t_)*. Note that this applies even when the sounds generated by the same source are separated by one or more sounds associated with different sources. The only transition that matters is that between the most recent tone and the last tone *in the same stream*, so if three tones *f_1_ f_2_* and *f_3_* had all been associated with the same stream, we would only consider the transition from *f_2_* to *f_3_*, whereas if *f_2_* was associated with a different stream, we would only consider the transition from *f_1_* to *f_3_*.

If a sound comes from a new source, then we assume that the likelihood is independent of previous tones:

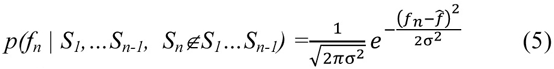

where 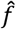 is the midpoint of the range of auditory frequencies presented for the trial. The final model has two parameters, *α* and *σ*.

### Posterior approximation

Using the iterative scheme above we can calculate analytically the possible combinations of tones, but as the tone sequence progresses the number of possible source combinations – and hence the size of the posterior distribution – increases exponentially. To prevent combinatorial explosion two methods are used to generate an approximation of the full posterior distribution. The first limits the number of tones that are retained when using the previous posterior as the next prior, i.e. the algorithm only retains e.g. the last 10 tones and their potential allocations to sources

Limiting the number of tones eases the computational load, and can also be seen as a crude model of a limited memory capacity. Although the iteratively constructed prior retains some stream information of all previous tones, when a very short memory is used this may not be sufficient to generate stable stream allocation as the CRP prior probabilities fluctuate greatly when the number of previous tones is small. Furthermore, if the structure of the sequence is an important cue for streaming, a larger memory may be necessary to determine regularities in the sequence.

Even when the memory is limited to the previous six tones, allocating a stream to the seventh tone requires a posterior distribution taking 858 values, most of which must necessarily have very small probabilities. A second method to limit the size of the posterior is simply to select only the most probable stream combinations by imposing a probability threshold, hence we only used stream combinations with *p*>0.001. Together these approximation methods allow a reasonable memory length of 10 tones (to avoid instability), while avoiding combinatorial explosion.

## Experiment 1

To compare the model to human performance we conducted a psychophysics experiment, in which six participants with normal hearing listened to simple auditory sequences and performed a subjective judgment task (a variant of experiments by van Noorden (1975)). Subjects were undergraduate students and received course credits for their participation. Each subject was fully briefed, provided informed consent and was given brief training on the task through exposure to 5 trial stimuli.

### Experimental setup

Figure 1a shows a schematic of the stimuli used – each sequence comprised 30 tones in repeated LHL-triplets, where the dash represents a silent gap. Each tone was 50 ms in duration, including 10 ms raised cosine onset and offset ramps. A 2×2 factorial design was used: the pitch of the high tones taking values of 3, 6, 9, 12 and 15 semitones above the low tone, which had a fixed frequency of 1000 Hz, and the offset to onset interval taking values 17, 33, 50 and 67 ms. The duration of the silent gap was equal to the tone duration plus the offset-onset interval. Conditions were ordered randomly – each condition was tested 20 times over 5 runs, each run lasting approximately 7 minutes. Stimuli were presented through Sennheiser 280 headphones at a comfortable supra-threshold level. At the end of the sequence participants pressed a key to report whether the percept at the end of the sequence was most like a single stream (a galloping rhythm) or two separate streams of notes. The fraction of 2-stream responses per condition is shown in figure 2b for all six participants.

**Figure 2:**
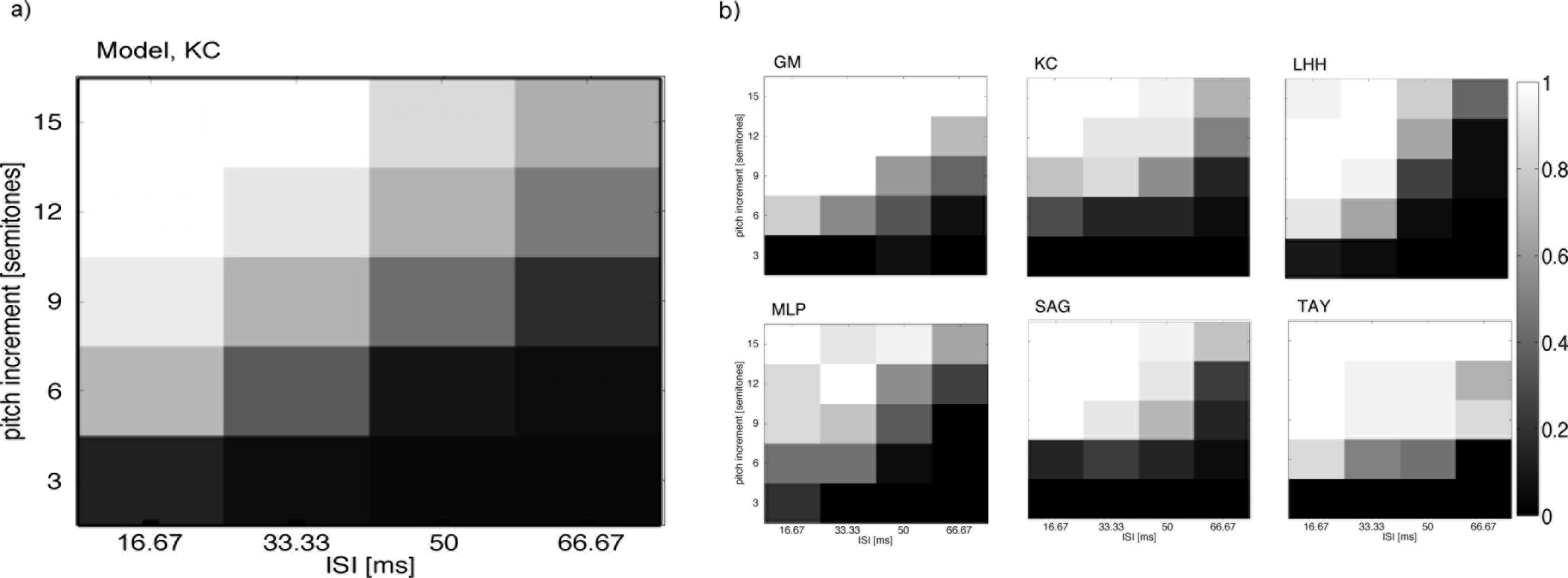
a) Model pred iction, based on fitted parameters from subject KC, giving the fraction of trials in which participant responded ‘2’ for the number of streams perceived. Axes give the pitch difference for the middle tone and the inter stimulus interval (ISI): the time between the offset of one tone and the onset of the next. b) The results from 6 subjects.

### Model response

To determine the response of the model to a tone sequence, the posterior for each possible sequence, *C*, is calculated tone-by-tone until all 30 tones have been presented. To relate the final posterior over sequences to subject responses, *s_r_* (‘1 or 2 streams’) *P_model_(s_r_|tones, C)*, we defined a metric between two sequences. While the simple Hamming distance was considered we found it did not capture the similarities and differences between sequences. As an example, the Hamming distance between the sequence [11111] and [12222], H(11111, 12222)=4, does not capture the intuition that a change of labels (2->1) implies a distance of 1. Instead we define a transition matrix, *M_C_* with elements *m_i, j_=C_i_-C_j_* i.e. the difference in the stream number for entry *i* and *j* of sequence *C*.

A transition matrix *M_pC_* is calculated for each posterior stream combination *C*, and also for the ‘ideal’ one or two stream response percepts (i.e. *M_1_* corresponding to 111 111… and *M_2_* corresponding to 121 121…). The sum of the absolute difference between elements of *M_pC_* and both *M_1_* and *M_2_*, *d_C1_=| M_pC_ – M_1_|* and *d_C2_=| M_pC_ - M_2_|* give measures of the distances *d_C1_* and *d_C2_* from *C* to the ideal response percepts. This method can also give the fixed distance d_12_ between the ideal responses, *d_12_=|M_1_-M_2_|*, thus streams *C*, 111 111… and 121 121… are represented by a triangle with sides of length *d_C1_*, *d_C2_* and *d_12_*. The vertex corresponding to stream *C* can be projected onto the side *d_12_* giving *D_1_*, the relative difference between *C* and the two response percepts:

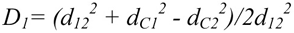

*D_1_* is restricted to the range [0, 1], and each projected point is weighted by its posterior probability to give the marginal distribution of the posterior projected onto the axis joining the two responses. The distance *D_1_* gives the probability of subjects response, *s_r_*, 1 or 2, given *C*, i.e. *P(s_r_ = 2|C)* = *D_1_* and *P(s_r_ =1|C) = 1-P(s_r_ =2|C).* Lastly we marginalize over the possible sequences, and assume that participants draw a sample from the posterior when responding, giving

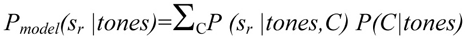

The parameters of the model (as well as for the alternative models below) were optimised using the MATLAB fminsearch routine to maximise the log-likelihood of the data, Σ *ln(P_model_(s_r_|tones))* independently for each subject. During each iteration of the search, a sequence of 30 tones was presented to the model for each condition, and the probability of response ‘1’ was calculated per condition.

### Model performance and comparison

The model was compared against three alternatives that used different priors to constrain the number of possible streams to two:

A. When the stream combination comprised only one stream (repeated), the prior probability of the next stream being 1 or 2 was allocated according to the CRP, but if the combination already contained two streams, the prior probability of allocating stream 1 or 2 was simply the fraction of previous tones that were allocated to stream 1 or 2 respectively.
B. The prior probabilities of a new tone being allocated to stream 1 or stream 2 was given by *P_1_*, and *1-P_1_* respectively, where *P_1_* is a free parameter.
C. The prior probabilities of a new tone being allocated to stream 1 or 2 were fixed at 0.5.

As mentioned earlier, an alternative response measure based on the Hamming distance was also tested: in this case we used the original, unconstrained CRP prior model. In the results, this is referred to as alternative D.

Because alternative model C has only one free parameter (all others have two), we use the Bayesian information criterion (BIC=*-2log P(resp|tones)*+*k*\**log(n)*, where *k* is the number of parameters and *n* is the number of data points fitted over) to compare model performance in table 1. With the exception of participant LHH, the unconstrained model gives a better fit (smaller BIC) than all the alternatives considered. The mean ± SEM of the optimised parameters for the unconstrained model are *α* = 0.81 ± 0.12 (equivalently *P(11)* = 0.56 ± 0.04) and *σ*= 105 ± 7 [semitones/sec]. Data from all subjects and the unconstrained model output for participant KC is shown in figure 2.

**Table 1.**
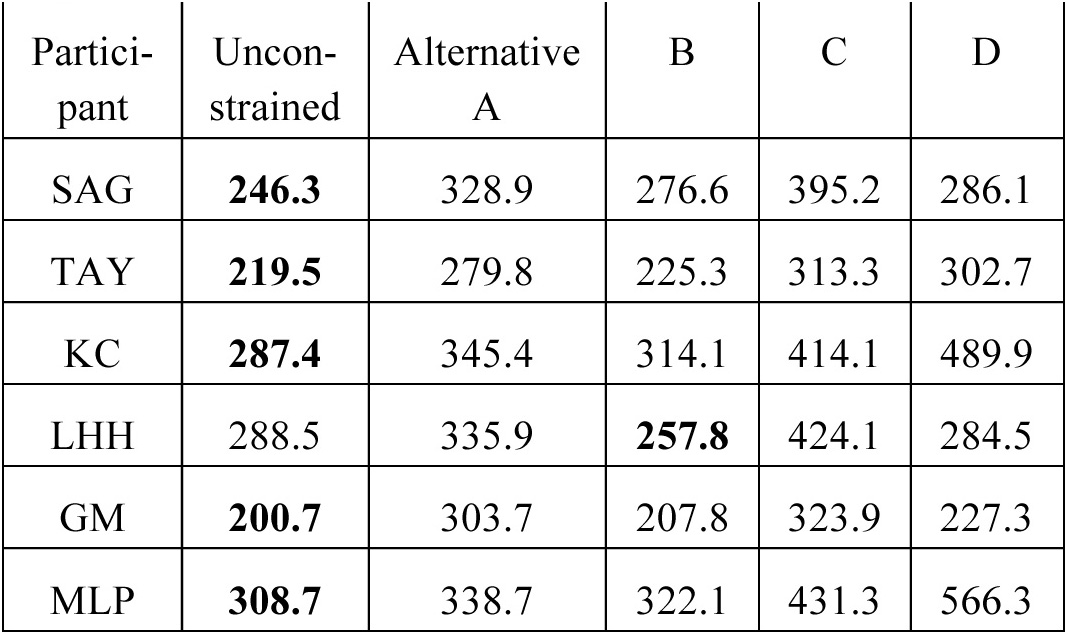
BIC per participant. A smaller number indicates better relative performance (best model for each subject indicated in **bold**).

| Parti-<br>cipant | Uncon-<br>strained | Alternative<br>A | B | C | D |
| --- | --- | --- | --- | --- | --- |
| SAG | **246.3** | 328.9 | 276.6 | 395.2 | 286.1 |
| TAY | **219.5** | 279.8 | 225.3 | 313.3 | 302.7 |
| KC | **287.4** | 345.4 | 314.1 | 414.1 | 489.9 |
| LHH | 288.5 | 335.9 | **257.8** | 424.1 | 284.5 |
| GM | **200.7** | 303.7 | 207.8 | 323.9 | 227.3 |
| MLP | **308.7** | 338.7 | 322.1 | 431.3 | 566.3 |

## Experiment 2

While the model above theoretically allows an unlimited number of tones to be segregated into an unrestricted number of streams, the classical experiment (presented above) only allows a sequence of 3 tones to be separated into 1 or 2 streams. However, the model predicts that subjects should generally segregate based on frequency and temporal distances between tones. To test this further we performed a novel follow-up experiment where subjects were presented with seven tones and had to indicate the number of streams perceived. Nine conditions were created with sequentially larger discrepancy in frequency between tones and thus a larger probability of being assigned to different streams according to the model. The temporal gap between tones (ISI) were kept constant at 33.3 ms, unlike experiment 1. For each condition, of the seven tones (see fig 1b for one condition) three tones were unique. Five further subjects (see above) performed this new task. Results showed that subjects perceived an increasing number of streams (fig. 3), in accordance with predictions from the model, rising from 1 to close to 3 (p<0.0001, F=20.39, one-way anova, df=8). None of the subjects perceived more than 3 streams for any of the conditions.

**Figure 3:**
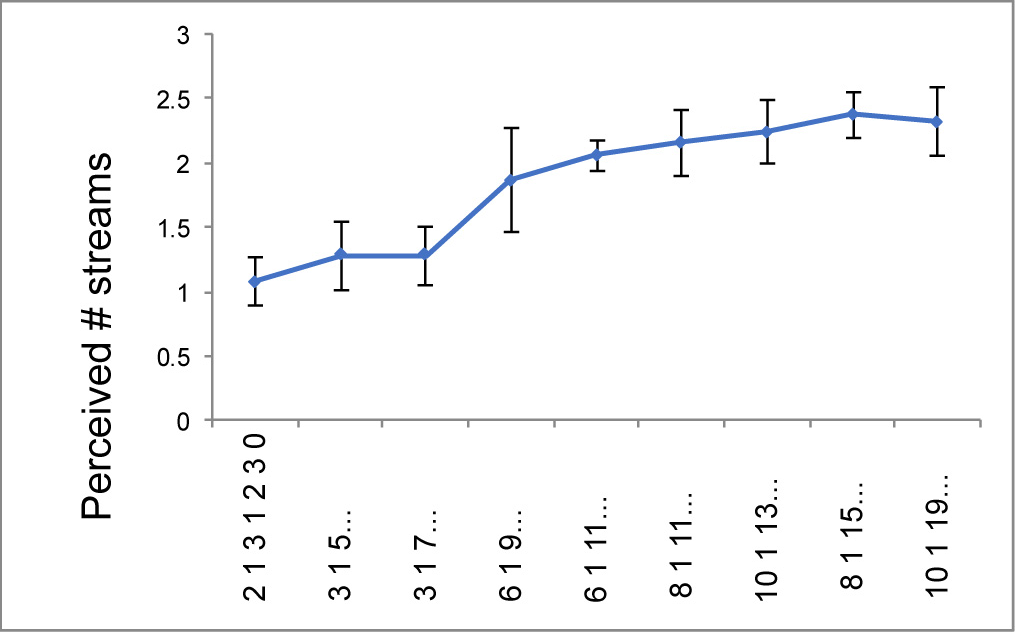
Averaged subject responses as a function of the auditory tone condition (see example in Fig. 1b). The horizontal labels indicate the tone-sequence of the condition, ordered by increasing step sizes. Error bars are standard errors.

## Discussion

We have presented a simple Bayesian statistical model for grouping of discrete sequential stimuli. Utilizing a non-parametric Bayesian prior the model iteratively updates the posterior distribution over the assigned group of each stimuli and provides an excellent description of the perceptual interpretation of simple auditory sequences in human observers.

With just two parameters, the model gives a good account of the basic characteristics of auditory stream segregation – the variation in the probability of perceiving a single sound source as a function of the repetition rate and pitch difference of the sounds. Although the ultimate goal is to segregation, for experimental simplicity we tested a well known paradigm from auditory psychophysics. The proposed model gave a better fit to the data than alternative models that were constrained to interpret the sounds as being produced from just one or two streams. Predictions from the model were also in accordance with results from a novel experiment with larger number of tones (exp. 2).

Importantly the model goes beyond giving just the number of sources, but says which sounds are produced by each source. While the combinatorial space of the posterior distribution in experiment 1 was collapsed to give a marginal distribution in a continuous 1-d response space (leading to an estimate of response probability), the maximum a posterior (MAP) for all participants was always located at either 111-111… or 121-121…, depending on the stimulus condition (figure 2b). This is reassuring as it is consistent with the anecdotal evidence that participants always perceive either a galloping rhythm (streams 111-111…) or a high-pitch and a low pitch stream (121-121…), i.e. the percept is always at the MAP. Indeed, the percept cannot in general be at the mean because the space of possible percepts is discrete: there is no percept between, say, 111 and 121.

One consequence of the inference model that is not addressed by mechanistic models of stream segregation is that when a percept changes from say 111-111 to 121-121, the source allocation of *previous* sounds is changed. Ironically, this ‘non-causal’ effect is essentially a feature of causal inference – when an observer decides that the percept has changed to 121-121, this is based on previous evidence, and yet at the time that the previous tones were heard, they were all associated with one source. A similar effect is commonly encountered when mis-interpreted speech (perhaps mis-heard due to background noise) suddenly makes sense when an essential word is heard – the previous words are reinterpreted, similar to the letters in predictive text message systems.

The framework of the model is very general, and allows for the incorporation of other factors into the likelihood to describe other aspects of auditory stream segregation. Adding terms in the likelihood function may be able to explain other effects seen in the literature, such as segregation based on bandwidth (Cusack & Roberts, 2000), or build-up and resetting of segregation (Roberts, Glasberg, & Moore, 2008). Furthermore, in the current study we assume that there is no ambiguity in the percept of the pure tones, the uncertainty arises from lack of knowledge about the underlying generative structure of the data. In a realistic situation perceptual ambiguity would have to be taken into account using an approach such as suggested by Turner and Sahani (Turner & Sahani, 2011). Nevertheless, we should emphasize that even though we are dealing with a Markov property (each tone within a stream only depends on the previous tone), the mixture of streams makes the problem very different from work on e.g. Hidden Markov Models (or even Infinite Hidden Markov Models) for which the goal would be to infer underlying states despite perceptual developed to separate audio signals (e.g. Roweis, 2001), these are not meant to mimic human perception, although a future comparison would be very interesting.

In the current implementation we used numerical approximations in order to handle the complexity of the model. As an alternative to calculating our results analytically we could use Monte Carlo techniques (e.g. Markov Chain Monte Carlo sampling, a different type of approximation), which have become a standard tool for solving complex statistical models.

The proposed model of auditory stream segregation is a specific instantiation of an iterative probabilistic approach towards inference of perceptual information. A major issue for this approach is the problem of dealing with multiple sources, as represented by the work done on causal inference (Shams & Beierholm, 2010). Until now models of causal inference have been unable to handle more than two sources, due to the escalating number of parameters needed for parametric priors. The use of a non-parametric prior allows a complex of many stimuli to be interpreted without running into this problem, potentially allowing for an arbitrary number of causes in the world. This approach is very general – it can be applied to any set of discrete sequential cues involving multiple sources – and it gives a simple, principled way to incorporate natural signal constraints into the generative model.

